# Rapastinel Accelerates Loss of Withdrawal Signs after Repeated Morphine and Blunts Relapse to Conditioned Place Preference

**DOI:** 10.1101/2022.08.19.504553

**Authors:** Christopher Armstrong, Julia Ferrante, Nidesh Lamichhane, C Gamard, Zachery Reavis, Q.D. Walker, N Zucker, A Patkar, Cynthia Kuhn

**Author notes:** Send correspondence to: Dr. Cynthia Kuhn, Box 3813 Duke university Medical Center, Durham NC 27710. Declaration of interest: Drs. Kuhn and Patkar have a patent pending on use of rapastinel in treating drug dependence. The other authors have no interests to declare.

## Abstract

The purpose of the present study was to evaluate the efficacy of rapastinel, an allosteric modulator of NMDA receptor function, to accelerate the loss of opioid withdrawal symptoms and blunt or prevent relapse to morphine conditioned place preference (CPP) in rats. Two studies were conducted. In study 1, adult and adolescent male and female rats were treated with increasing doses of morphine (5 mg/kg, bid to 25 mg/kg bid) for 5 days. On day 6 animals were treated with naloxone (1 mg/kg) and withdrawal was assessed. They were then treated with saline or rapastinel (5 mg/kg) on days 6 and 8, and withdrawal assessed on day 9. Rapastinel treated animals exhibited significantly lower levels of withdrawal signs on day 9. No sex or age differences were observed. In Study 2, CPP for morphine was established in adult rats (males and females) by 4 daily pairings with saline and morphine (am/pm alternation). They were tested for CPP on day 5, and then treated with rapastinel (5 mg/kg) or saline daily on days 6-10 of extinction. On day 11 they received a final dose of rapastinel or saline followed by extinction. On day 12, animals received 1 mg/kg of morphine and were tested for relapse. Rapastinel did not affect extinction of CPP, but rapastinel-treated animals spent significantly less time in the previously morphine-paired side than saline-treated animals during the relapse trial. These findings of accelerated loss of withdrawal signs and blunted relapse to CPP suggest that rapastinel could provide an adjunctive therapy for opioid dependence during initiation of pharmacotherapy for opioid dependence.

## Introduction

The current opioid crisis has generated demand for additional therapeutic tools for treating opioid dependent patients. Treatment with the opioid drugs remains the current mainstay of pharmacotherapy of opioid dependence at the current time, but there are significant limitations to each of the FDA-approved pharmacotherapies. The opioid agonist methadone improves clinical outcomes but maintains the opioid dependent state.^1, 2^ The partial agonist buprenorphine can elicit withdrawal signs if patients use opioids during treatment and the opioid antagonist naltrexone requires an opioid-abstinent period before treatment begins.^3^ Therefore, there is a pressing need for a non-opioid treatment to complement available pharmacotherapies. This need is especially critical for a rising population of adolescent opioid users. Adolescents and young adults (ages 18-25 especially) exhibit significant use of prescription opioids (14% of high school seniors reported use in the 2017 Youth Risk Behavior Study) and this demographic has experienced an alarming increased in opioid overdose deaths, likely due to the presence of fentanyl and other opioids in the opioids they consumed.^4^ Furthermore, there is only one medication approved for use in opioid-dependent adolescents (buprenorphine) which is effective but retention of adolescents in buprenorphine treatment has been poor and they show a high relapse rate.^5-7^

The role of glutamate neurons in the acquisition, expression and relapse to opioid reward has generated interest in glutamate receptors as potential pharmacologic targets to treat opioid dependence. Many aspects of glutamate transmission change during addiction, including glutamate release and uptake as well as both function of AMPA and NMDA receptors.^8-12^ Upregulation of NMDA receptor subunits (NR1) perseveres for weeks after chronic opioid exposure.^13-17^ This neuroadaptation of NMDA receptors may be critical for the development of compulsive use of opioids.^18-20^ Research on the ability of NMDA receptor antagonists to treat opioid dependence support this possibility.^21^ In animal models, the prototype NMDA antagonist ketamine diminished heroin, methamphetamine and cocaine self-administration,^22^ the NMDA antagonist MK-801 blunted opioid withdrawal,^23^ and the NMDA antagonist memantine reduced opioid withdrawal symptoms in one human trial.^24^ Unfortunately, each of these pharmacologic approaches has significant limitations. Ketamine has a known human abuse and a serious side-effect profile, provoking dissociation and hallucinations at high doses.^25^ Memantine is a weak antagonist, and its clinical action is limited. Subtle modulation of NMDA receptor function has also been explored.^26^ D-cycloserine, a glycine-site partial agonist, facilitated extinction of morphine CPP as well as naloxone conditioned place aversion,^27-29^ and another glycine-site antagonist felbamate diminished withdrawal severity in animals.^30^ However, d-cycloserine has been unsuccessful in clinical studies, which may reflect narrow dose range, short time course and lack of efficacy with chronic treatment.^31, 32^

The purpose of the present study was to investigate the possibility that rapastinel, a novel allosteric modulator of the NMDA receptor, might show efficacy in treating opioid dependence without the limitations that other glutamate-targeted drugs have demonstrated. Rapastinel is a brain penetrant tetrapeptide derived from a monoclonal antibody to the NMDA receptor that functions as a partial agonist at the glycine site of this receptor.^33-36^ It has shown potential efficacy against depression and PTSD, reversal of cognitive impairment and neuroprotection without the psychiatric side effects or reinforcement actions of ketamine.^35, 37-40^ Rapastinel may act in part through enhancement of NMDA receptor NR2B subunit action.^34, 41-43^ We hypothesized that rapastinel could suppress withdrawal signs, perhaps prevent or blunt relapse and treat well-known co-morbidities like depression that are a barrier to recovery.

The present study included both males and females to determine if there was a sex-specific sensitivity to treatment, as well as both adolescents and adults.^5-7^ Successful trial data with N-acetylcysteine against cannabis suggest that glutamate-targeted therapy may be effective in this undertreated population.^44^

To test this hypothesis, we this study utilized experimental designs that were intended to model treatment initiated at the beginning of abstinence, a situation analogous to the one that patients would experience. In experiment 1, we monitored opioid withdrawal symptoms with a well-characterized investigator-administered morphine treatment paradigm followed by assessment of naloxone-precipitated withdrawal at two time intervals. Rapastinel treatment was initiated at the start of opioid *abstinence*, after the first naloxone-precipitated withdrawal, to mimic clinical experience. Animals were treated with rapastinel or saline for three days, and naloxone-precipitated withdrawal was repeated to determine if rapastinel accelerated the loss of opioid dependence. The timing of the trial was based on substantial literature demonstrating that passive opioid withdrawal symptoms persist for at least a week in rats after a brief chronic treatment regimen.^45-47^ The second experiment evaluated the ability of rapastinel to blunt relapse after a brief (5 day) conditioned place preference paradigm. Rapastinel treatment was initiated during extinction from CPP, again paralleling potential clinical practice, and its effect on relapse after a probe dose of morphine was tested. A brief conditioning period and low dose of morphine were utilized to achieve extinction over a brief period. This facilitated timely assessment of rapastinel efficacy.

## 1. Materials and Methods

### Experimental Subjects

Subjects were male and female adolescent and adult rats (PN 28-30 and PN 70-72) from Charles River Laboratories (Raleigh NC). Animals were grouped-housed in single sex groups in a 12/12 light/dark cycle with free access to water and standard lab chow.

### Drug Treatments

The basic treatment regimen is shown in **Figure 1**. Rats were treated with a 5-day, increasing dose morphine regimen (5 mg/kg sc, bid, increasing 5 mg/kg/day to 25 mg/kg). Controls were treated with sterile 0.9% saline bid. Adult male and female rats were randomized to saline or rapastinel post-morphine treatment groups before initiation of morphine treatment. All animals received saline or morphine on days 1-5, and a dose of naloxone (1 mg/kg, sc) on the morning of day 6 (12 hours after the last morphine dose) to precipitate withdrawal. Withdrawal behaviors (wet dog shakes, diarrhea, mastication, salivation, ptosis and abnormal posture) were quantified as described by Gellert and Holtzman^48^ starting 10 min. after naloxone. They were then treated with saline or rapastinel (5 mg/kg on day 6 and 8) according to group designation. The dose and alternate days of treatment are based on rat studies of rapastinel effects on depressive-like behavior which persisted for several days after a single injection.^41, 49^ On day10, animals received a second naloxone challenge (1 mg/kg, sc) and withdrawal was scored again. One female was removed from the withdrawal cohort due to values triple that of other animals (excluded by Grubbs test).

**Figure 1.**
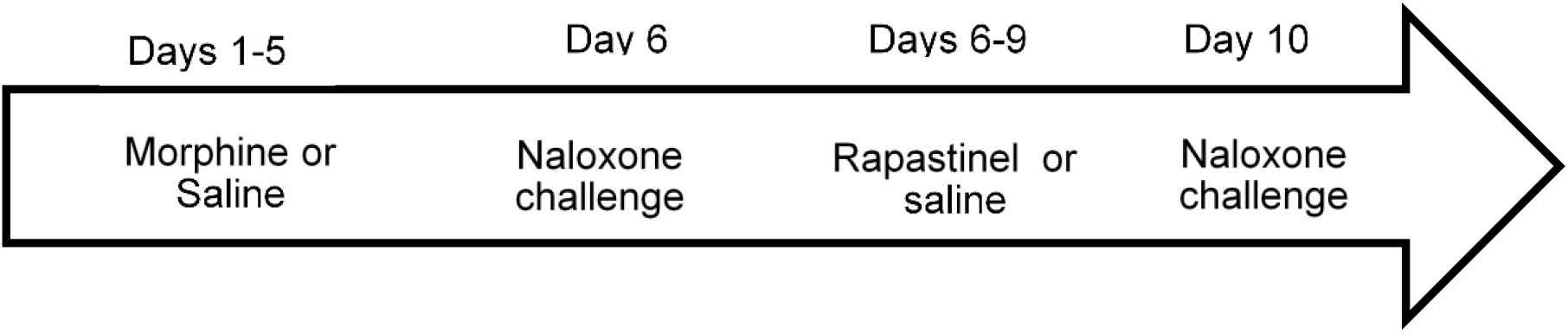
Experiment 1.

### CPP and extinction

Conditioned Place Preference (CPP) was conducted in adult male and female rats with a biased CPP procedure adapted from Mueller et al^50^ (Figure 2). CPP was conducted in a two-sided chamber. Side 1 had white walls and plexiglass flooring, and Side 2 had black walls and plastic mesh flooring (preferred side). <u>Acquisition</u>: On habituation day (day 1), animals were allowed to explore the whole apparatus for 30 minutes. On days 2-4, rats injected with morphine (2.5 mg/kg sc) or saline and immediately restricted to Side 1 for morphine or Side 2 for saline (alternated am and pm daily). On test day 5, time on each side was monitored as animals explored the apparatus freely. <u>Extinction</u> was conducted on days 6-10. Rats received saline pairings in both sides once daily. During the extinction trial on day 10, animals could explore the entire chamber. <u>Relapse</u> was measured on day 11: animals were treated with saline or morphine (1 mg/kg) and allowed to explore both sides of the apparatus. Animals were assigned to saline or rapastinel before the experiment started but were treated with saline or rapastinel (5 mg/kg) only during extinction in the morning on each day. Data are expressed as time on the drug-paired side. Four animals from each group were excluded due to failure to develop a CPP and so extinction and relapse were not tested.

**Figure 2.**
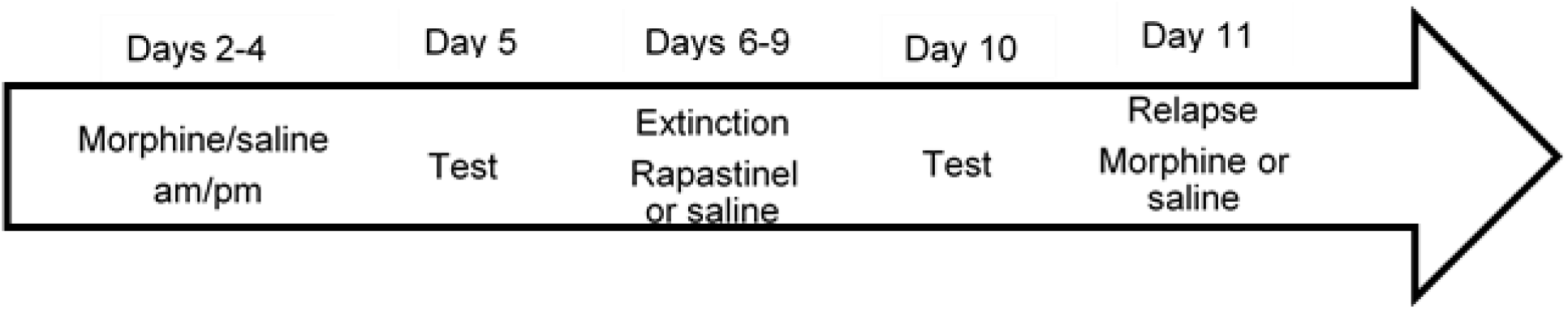
Experiment 2.

### Materials

Morphine was supplied by the National Institute for Drug Abuse. Naloxone was obtained from Sigma Chemical Company (St Louis, MO). Rapastinel was synthesized by the Duke Small Molecule Synthesis Facility and provided by Dr. David Gooden.

### Statistics

Withdrawal results were analyzed by 3-way repeated measures ANOVA (age, sex and treatment was the between subject variables and day was the within subject variable) followed by post-hoc Fishers LSD multiple comparison test to compare differences between groups. CPP data were analyzed by repeated measures ANOVA (treatment and sex the between subject variables and day as within-subject variable). All experiments were approved by the Duke University IACUC and conducted in accordance with the NIH Guide for the Care and Use of Laboratory Animals.

## 2. Results

Figure 3. shows mean withdrawal scores of male and female adults on Day 6, and on day 9, after 2 treatments with saline or rapastinel. Day 6 data are shown with combined saline-rapastinel groups as these data were collected before cohorts were treated. Animals exhibited robust withdrawal signs which diminished by day (main effect of day, F = 122.2, p < 0.0001 for effect of day). Animals treated with rapastinel had significantly lower withdrawal scores on day 9 (Treatment x day interaction, F = 17.95, p < 0.0004). There was a sex x day interaction (F = 12.00, p < 0.002) but no interaction with treatment: males declined more than females from day 6 to day 9. Responses in adolescents were similar to those observed in adults.

**Figure 3.**
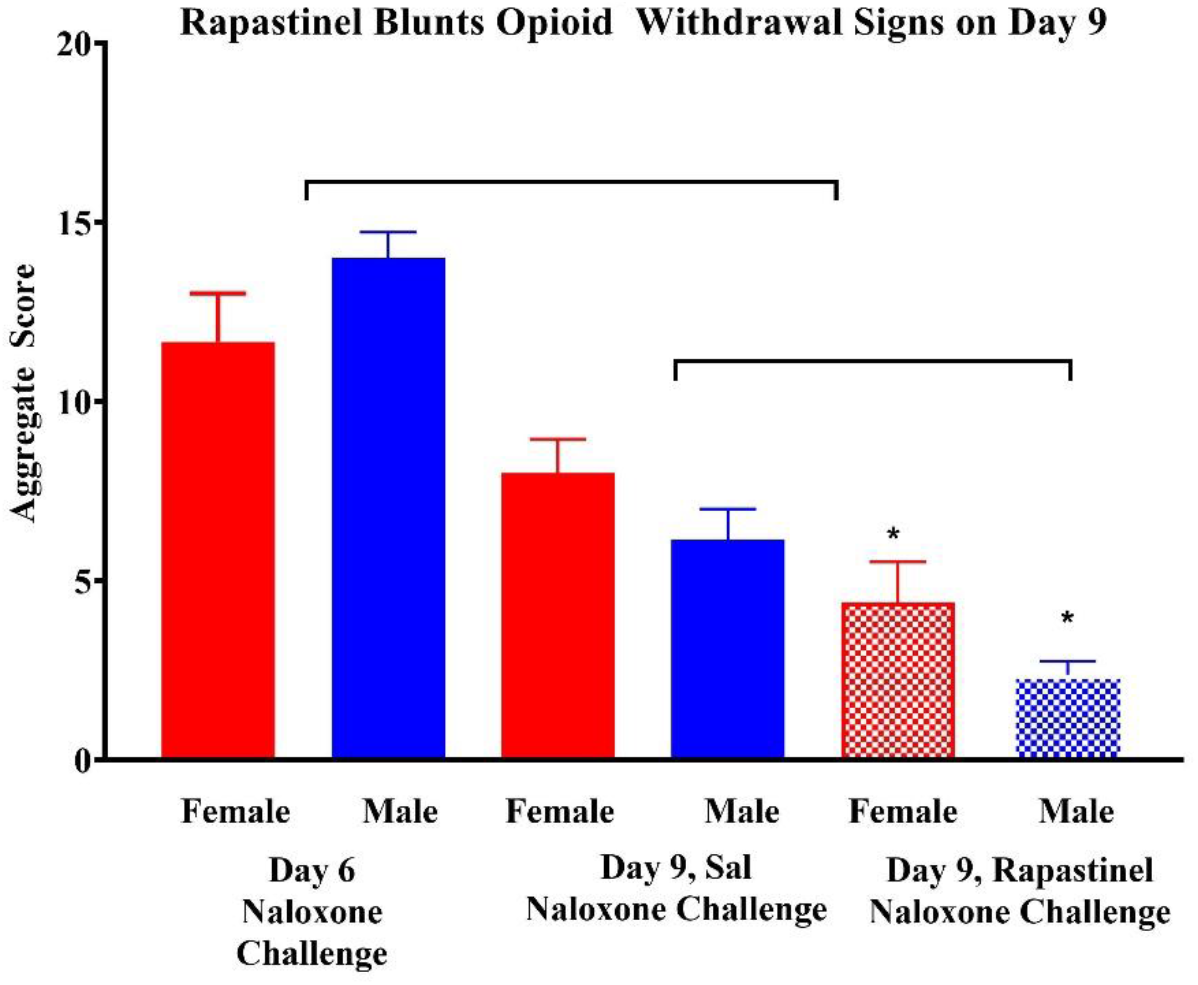
Naloxone-precipitated withdrawal score on days 6 and 9 after 5 days of morphine treatment. Females indicated in red, males in blue. Rapastinel treatment indicated as cross-hatched. Results expressed as mean ± SEM. Brackets indicate day 6 different from day 9 and Day 9 Saline different from Day 9 Rapastinel by Fisher’s LSD multiple comparison test. N’s 5-7 for each experimental group.

Figure 4. shows withdrawal on Day 9 after post-morphine saline or rapastinel in adult rats (adult data from Figure 1) and a similar cohort of adolescents. Results are collapsed for sex, as no main effect of sex or interaction with treatment was observed. N’s for the groups collapsed by sex were 12-14/group. Withdrawal on Day 6 was comparable in adolescents and adults (adult = 13.5 ± 0.9, N = 25, adolescent = 12.3 ± 0.7, N = 28). Naloxone treatment elicited a significant withdrawal response which declined with days after the end of morphine treatment (F =201.9, p < 0.0001 for effect of day). Animals treated with rapastinel had a lower response on day 9 compared to saline-treated animals (F = 31.1, p < 0.0001 for interaction of day x treatment, testing on day 6 vs. day 9), suggesting that treatment with rapastinel accelerated the loss of dependence over this brief time period in both adolescents and adults. There was no main effect of age or interaction or age x treatment.

**Figure 4.**
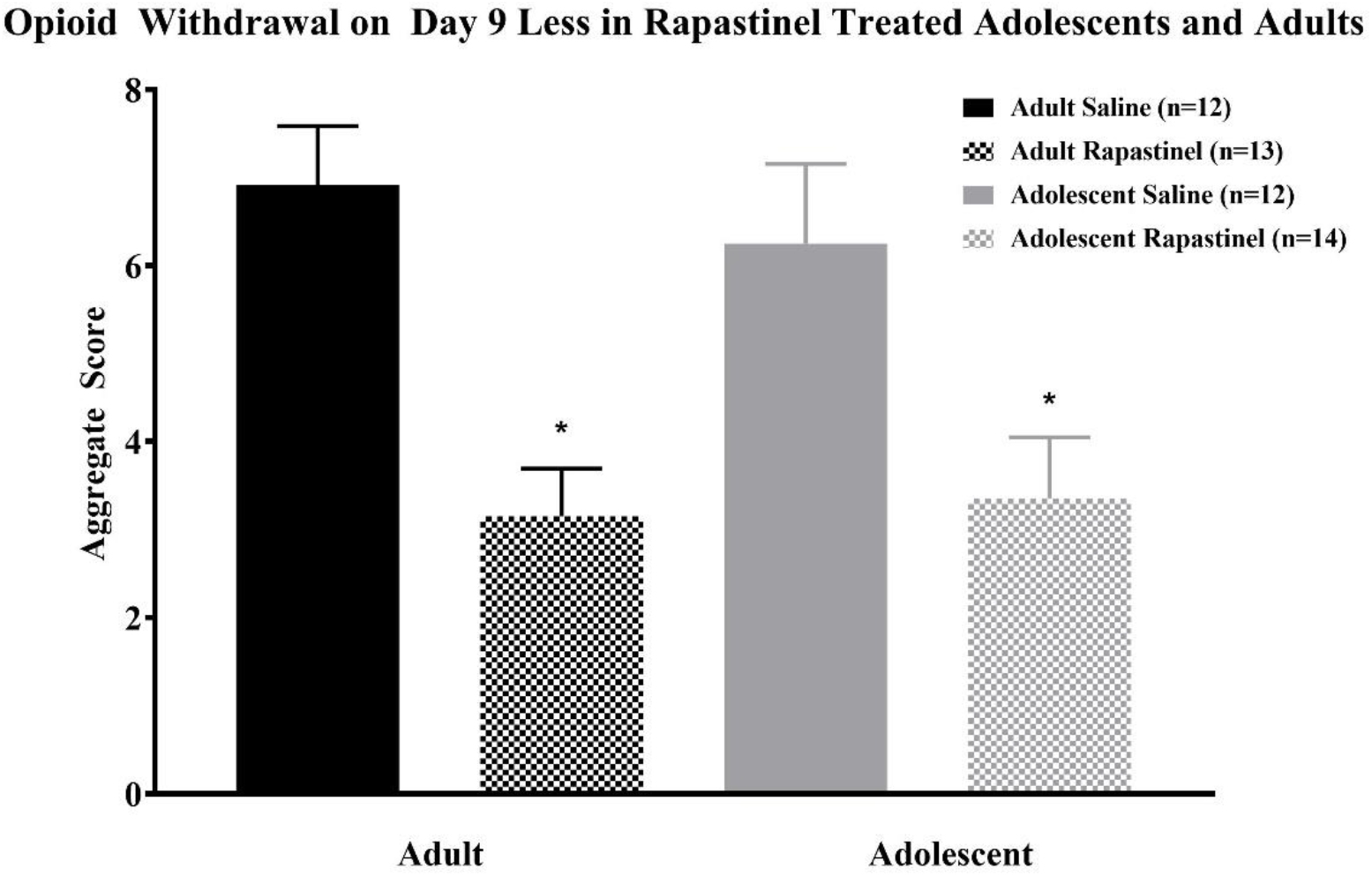
Naloxone-precipitated withdrawal on Day 9 in adults and adolescents treated with saline or rapastinel from day 6 to day 9 after a 5-day morphine treatment. Results are expressed as mean ± SEM. * indicates different from saline treatment by Fisher’s LSD multiple comparison test. + indicates different from age-matched Saline. N’s for the groups collapsed by sex were 12-14/group.

CPP was conducted only in adults, as no age differences were observed in the withdrawal study. Repeated measures ANOVA including sex and treatment as between factors and time (1 = habituation, 2 = CPP and 3 = extinction) as within factors. Rapastinel was included as a treatment to indicate that there was no difference between animals assigned to saline or rapastinel BEFORE treatment started.

**Figure 5a** shows habituation, CPP to morphine and Extinction in males and females. The repeated measures ANOVA indicated a significant effect of time indicating significant increase in time spent in the drug-paired side followed by decrease during extinction (F (1,34) = 55.5, p < 0.0001, N = 18 for saline, 17 for rapastinel). Time 1 (habituation) was significantly different from CPP and CPP was significantly different from Extinction by Fishers LSD multiple comparison test. No sex differences were observed, and no effects of rapastinel (as expected, as rapastinel treatment was initiated after CPP was established, and expected only in Time 3 (Extinction).

**Figure 5.**
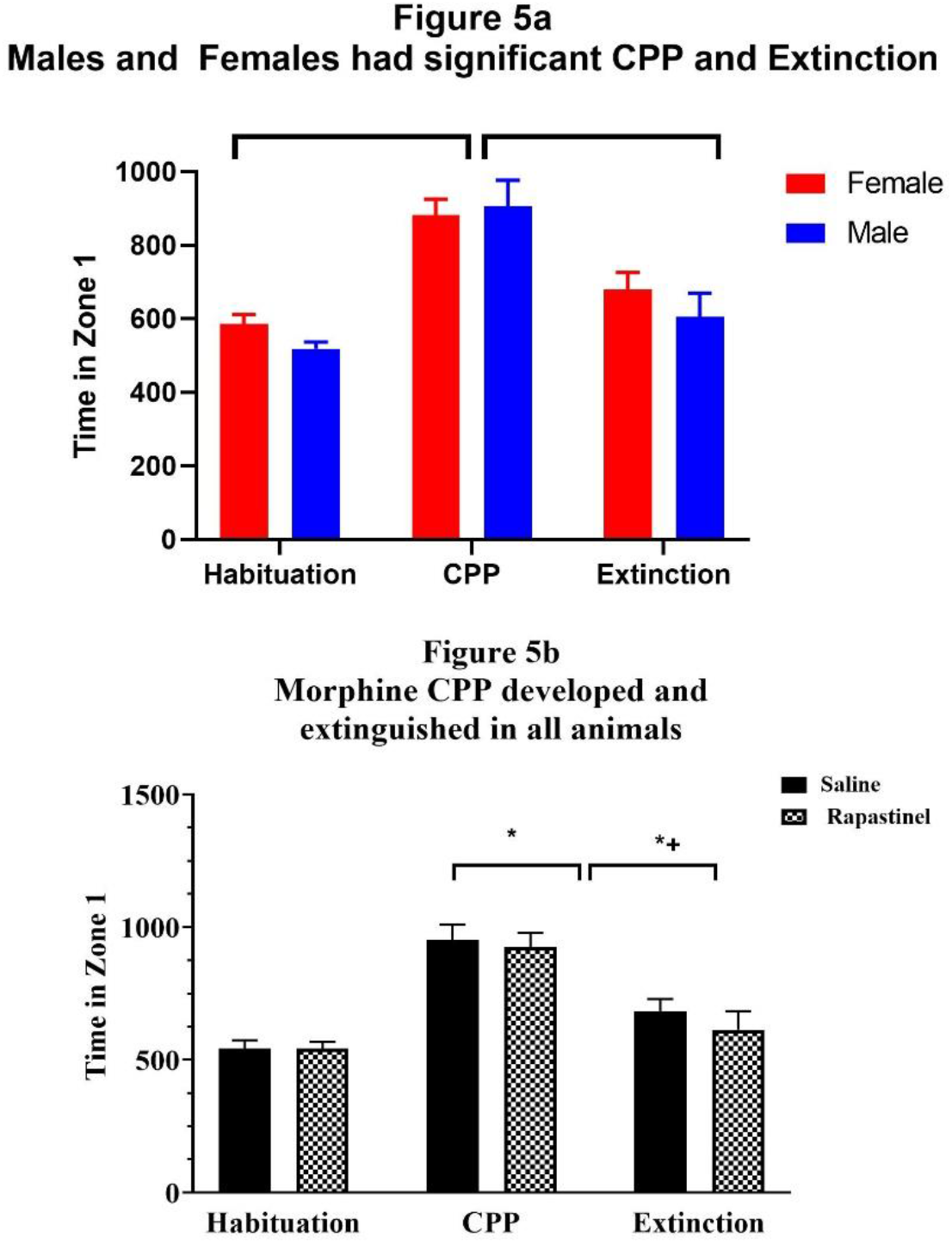
a. Habituation, CPP and Extinction during morphine-induced CPP by sex. Results are expressed as mean ± SEM. Brackets indicate CPP different from Habituation and from Extinction by Fishers LSD multiple comparison test. N = 19 for females and 16 for males. B. Results collapsed by treatment. Saline-treated animals are solid, rapastinel-treated animals are cross-hatched. * indicates different from habituation and + indicates different from rapastinel by Fishers LSD multiple comparison test. N = 18 for saline, 17 for rapastinel).

**Figure 5b** shows data by treatment (pooled by sex) to demonstrate that there was no difference in habituation or CPP times for animals pre-assigned to saline or rapastinel groups (treatment which began after assessment of CPP through extinction, indicating no baseline differences in the magnitude of CPP. Extinction values were slightly but significantly still elevated relative to habituation.

Figure 6. shows extinction and relapse data collected after all animals were treated with morphine following extinction as shown in Figure 2. Repeated measures ANOVA was conducted with sex and treatment was between factors, and time (time 1 = extinction, time 2 = relapse) as within factors, as indicated above. ANOVA indicated a significant effect of day (F = (1,33) = 9.97, p < 0.003), a significant effect of treatment and a significant interaction of time x treatment interaction (F 1, 33) = 6.55, P < 0.02) indicating that rapastinel blunted relapse. There was no effect of sex on any of the measures.

**Figure 6.**
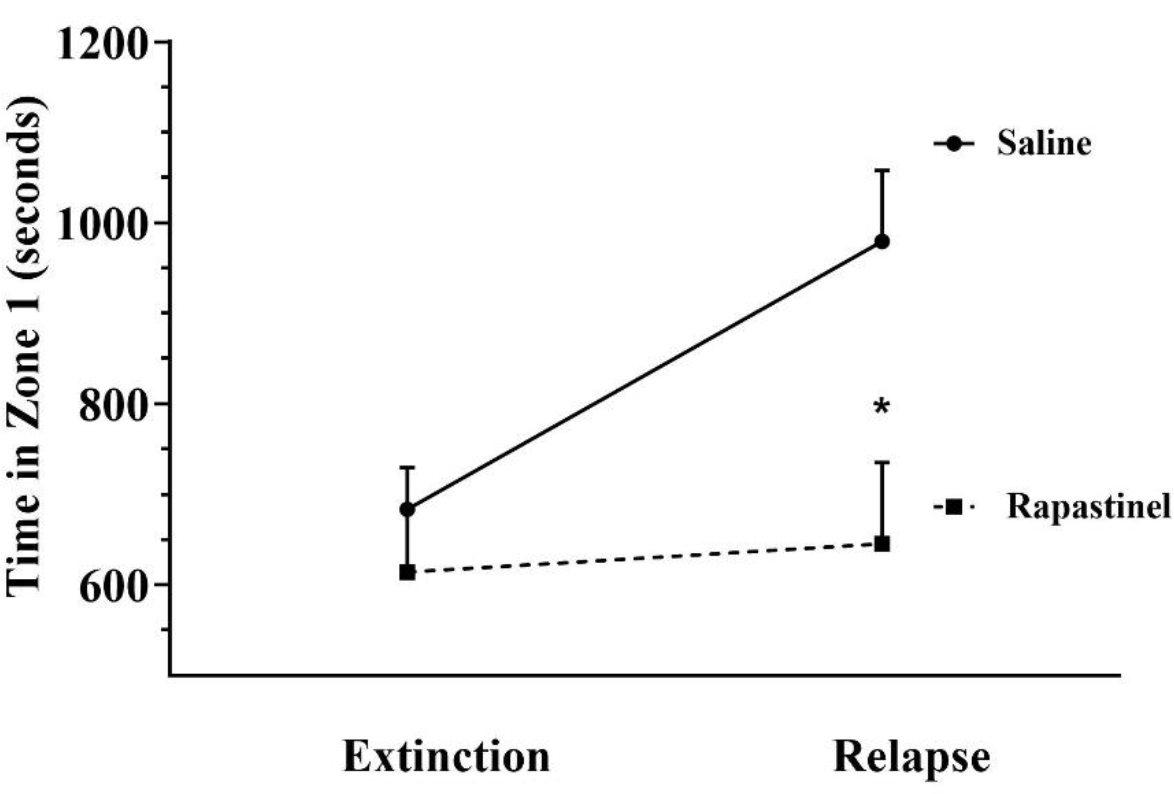
Rapastinel Prevented Relapse to Opioid Conditioned Place Preference. Relapse to CPP with morphine challenge in animals treated with saline (solid line) or rapastinel (dotted line. Results are expressed as mean ± SEM. * indicates different from saline treatment by Fisher’s LSD multiple comparison test. N = 18 for saline and 17 for rapastinel).

## 3. Discussion

The main findings of this study are that rapastinel, the novel allosteric modulator of NMDA receptors for glutamate rapastinel can accelerate the loss of opioid dependence in animals that have become morphine-dependent through investigator-administered drug and can blunt relapse to morphine conditioned-place preference. Rapastinel treatment initiated at the start of withdrawal (as would occur for a clinically useful pharmacotherapy) resulted in lower withdrawal signs on day 9 in both sexes and ages and treatment begun during extinction of CPP resulted in blunted relapse to morphine CPP. We speculate that these changes may represent a more rapid reversal of neuroplastic changes that contribute to opioid dependence. These two findings suggest that it could potentially be useful in treatment of opioid dependence.

The rapastinel dosing strategies utilized in this study were based on two considerations. First, previous reports demonstrated that rapastinel has a brief half-life within the body, but prolonged biological/behavioral effects after a single dose. This has been demonstrated in numerous studies of its effect on pain, and in animal models of depression.^36, 37, 51, 52^ Second, we initiated treatment *after* the development of dependence or CPP to evaluate its potential therapeutic utility without interference from its potential to blunt the development of dependence or place preference. A slightly different regimen (daily not alternate days) was chosen for CPP study to guarantee coverage of both extinction and relapse in the absence of any previous research on this agent in this behavioral protocol. This approach models how rapastinel might be used in a clinical context, in which patients are already drug dependent, but could benefit from treatments which ameliorate withdrawal or lower the chance of relapse.

All animals, regardless of sex or age, demonstrated marked and comparable signs of naloxone precipitated withdrawal twelve hours after the last morphine treatment, and smaller but still statistically significant signs on day 9, after 3 days of passive withdrawal. The fall in withdrawal signs from day 6 to day 9 is consistent with an abundant literature on opioid withdrawal in adults which shows that withdrawal signs are typically observed for up to 8-10 days after cessation of a treatment regimen.^53-55^

The lack of sex or age differences in the magnitude of withdrawal could reflect the increasing dose regimen of morphine and the large dose of naloxone used which resulted in high withdrawal scores. Exaggerated naloxone-precipitated withdrawal has been reported in males compared to females^56^ as has the absence of sex differences.^54, 57^ Opioid withdrawal occurs even in infancy in humans^58^ and rats,^59, 60^ but behavioral manifestations differ and mechanisms differ from adults.^61^ We showed that behavioral and hormonal withdrawal responses increased from postnatal day (PN) 10 to PN 26 (juvenile, pre-adolescent age).^62^ The data comparable to adults is consistent with the increasing pattern observed in our previous study.^62^ There are few other studies of opioid dependence in adolescent rats and these involve other dependent measures, and no comparison to adults.^63^ The one study using comparable behavioral measures surprisingly reported no withdrawal signs in adolescents after heroin exposure,^64^ a finding that contradicts the literature reporting withdrawal signs at considerably earlier ages.

The lack of sex differences in morphine CPP is consistent with several studies which reported comparable morphine CPP in male and female rats at the dose used in the present study, although females have been reported to acquire CPP at lower doses than males.^65-67^ These CPP findings are consistent with the conflicting literature about sex differences in opioid self-administration and dependence in rodents. Sex differences (with females acquiring faster and taking more drug) have been reported^57, 68, 69^ as has the absence of sex differences.^70, 71^

Withdrawal signs decreased significantly more in rapastinel-treated animals than in saline-treated animals from day 6 to day 9, although the distribution of specific withdrawal signs did not differ (data not shown). The physiologic basis of the signs tracked by Gellert and Holtzman (wet-dog shakes, diarrhea, mastication, salivation, ptosis and abnormal posture) are skewed toward peripheral autonomic effects likely mediated by increased activation of descending noradrenergic function by afferent input.^72-75^ The effects of rapastinel observed in the present study are consistent with studies by Rasmussen which have shown that NMDA antagonists may blunt physical signs of withdrawal through actions in the locus coeruleus.^75-78^ The findings of these studies that ketamine-like behaviors limited the potential utility of targeting this mechanism to blunt opioid withdrawal illustrate the potential benefit of rapastinel, which lacks these effects.^37, 79^

Rapastinel also significantly decreased relapse to CPP following extinction, which likely was mediated by neural circuits distinct from those mediating the naloxone-precipitated withdrawal signs. The dose of morphine used in CPP studies was considerably lower than that used in the withdrawal study (4 daily doses of 2.5 mg/kg vs twice daily dosing increasing from 5 to 25 mg/kg/injection and animals experienced 5 days before the relapse study. Multiple previous studies show that NMDA antagonists memantine and MK801 blunt relapse to morphine CPP ^80-82^ although negative effects of d-cycloserine have been reported.^27^ Rapastinel values did not differ from controls after extinction in contrast with a recent study which showed that spermidine both enhanced extinction and prevented reinstatement in mice through an action the authors speculated reflected its modulatory action on NMDA receptors.^83^ However, extinction was complete in the present study before the test, while the latter study tested spermine effects before extinction was complete, Nevertheless, the successful blockade of relapse supports the potential clinical utility of rapastinel in preventing relapse in abstinent opioid dependent patients.

There has been significant research interest in the possibility that negative modulators of NMDA receptor function might prove clinically useful in treating opioid dependence, as outlined above. Many aspects of glutamate transmission change during addiction, including glutamate release and uptake as well as both expression and function of AMPA and NMDA receptors.^8-12^ Neuroadaptation of NMDA receptors may be critical for the development of compulsive use of opioids.^18-20^ Upregulation of NMDA receptor subunits (NR1) perseveres for weeks after chronic opioid exposure.^13-17^ The ability of rapastinel to blunt opioid withdrawal signs and relapse to CPP is consistent with studies of other NMDA receptor antagonists which blunt other behaviors reflective of opioid reinforcement. Rapastinel may serve as a first-in-class of a family of NMDA receptor modulators which lack ketamine-like side effects that are being explored for treatment of depression, pain and other conditions that may also prove useful in treating drug dependence. These include apimostinel, a more potent analogue of rapastinel and CP-101,606 (traxoprodil), which also targets NR2B receptors.^84, 85^

The present study utilized two rapid screens for pharmacologic action against opioid dependence (withdrawal, relapse to CPP). Future work is needed to clarify dose and frequency of rapastinel treatment needed to optimize these outcomes and utilize gold-standard approaches to studying opioid dependence including extended-access opioid self-administration. Investigation of its mechanism and location of action and assessment of its efficacy in blunting opioid-induced hyperalgesia and prolonged abstinence signs like “craving” and anxiety which are thought to play a significant role in relapse to opioid use will also provide substantial information about the utility of this therapeutic approach.^57, 86, 87^

## References

1. Inturrisi CE. Pharmacology of methadone and its isomers. Minerva Anestesiol. Jul-Aug 2005;71(7-8):435-7.

2. Inturrisi CE. Clinical pharmacology of opioids for pain. Clin J Pain. Jul-Aug 2002;18(4 Suppl):S3–13. doi:10.1097/00002508-200207001-00002

3. Maiti T, Das S, Ramasamy A, Stanley Xavier A, Kumar Behera S, Selvarajan S. An overview on medication-assisted treatment (MAT) for opioid dependence. J Opioid Manag. Mar/Apr 2020;16(2):141-149. doi:10.5055/jom.2020.0560

4. Bohm MK, Clayton HB. Nonmedical Use of Prescription Opioids, Heroin Use, Injection Drug Use, and Overdose Mortality in U.S. Adolescents. J Stud Alcohol Drugs. Jul 2020;81(4):484-488.

5. Marsch LA, Moore SK, Borodovsky JT, et al. A randomized controlled trial of buprenorphine taper duration among opioid-dependent adolescents and young adults. Addiction. Aug 2016;111(8):1406-15. doi:10.1111/add.13363

6. Marsch LA, Bickel WK, Badger GJ, et al. Comparison of pharmacological treatments for opioid-dependent adolescents: a randomized controlled trial. Arch Gen Psychiatry. Oct 2005;62(10):1157-64. doi:10.1001/archpsyc.62.10.1157

7. Schuman-Olivier Z, Weiss RD, Hoeppner BB, Borodovsky J, Albanese MJ. Emerging adult age status predicts poor buprenorphine treatment retention. J Subst Abuse Treat. Sep 2014;47(3):202-12. doi:10.1016/j.jsat.2014.04.006

8. Neuhofer D, Kalivas P. Metaplasticity at the addicted tetrapartite synapse: A common denominator of drug induced adaptations and potential treatment target for addiction. Neurobiol Learn Mem. Feb 9 2018;doi:10.1016/j.nlm.2018.02.007

9. Scofield MD, Heinsbroek JA, Gipson CD, et al. The Nucleus Accumbens: Mechanisms of Addiction across Drug Classes Reflect the Importance of Glutamate Homeostasis. Pharmacol Rev. Jul 2016;68(3):816-71. doi:10.1124/pr.116.012484

10. Medrano MC, Mendiguren A, Pineda J. Effect of ceftriaxone and topiramate treatments on naltrexone-precipitated morphine withdrawal and glutamate receptor desensitization in the rat locus coeruleus. Psychopharmacology (Berl). Aug 2015;232(15):2795-809. doi:10.1007/s00213-015-3913-2

11. Rawls SM, Baron DA, Kim J. beta-Lactam antibiotic inhibits development of morphine physical dependence in rats. Behav Pharmacol. Mar 2010;21(2):161–4. doi:10.1097/FBP.0b013e328337be10

12. Hearing M, Graziane N, Dong Y, Thomas MJ. Opioid and Psychostimulant Plasticity: Targeting Overlap in Nucleus Accumbens Glutamate Signaling. Trends Pharmacol Sci. Mar 2018;39(3):276–294. doi:10.1016/j.tips.2017.12.004

13. Anderson EM, Del Valle-Pinero AY, Suckow SK, Nolan TA, Neubert JK, Caudle RM. Morphine and MK-801 administration leads to alternative N-methyl-D-aspartate receptor 1 splicing and associated changes in reward seeking behavior and nociception on an operant orofacial assay. Neuroscience. Jul 12 2012;214:14-27. doi:10.1016/j.neuroscience.2012.04.032

14. Anderson EM, Reeves T, Kapernaros K, Neubert JK, Caudle RM. Phosphorylation of the N-methyl-d-aspartate receptor is increased in the nucleus accumbens during both acute and extended morphine withdrawal. J Pharmacol Exp Ther. Dec 2015;355(3):496–505. doi:10.1124/jpet.115.227629

15. Bajo M, Crawford EF, Roberto M, Madamba SG, Siggins GR. Chronic morphine treatment alters expression of N-methyl-D-aspartate receptor subunits in the extended amygdala. J Neurosci Res. Mar 2006;83(4):532–7. doi:10.1002/jnr.20756

16. Oh S, Kim JI, Chung MW, Ho IK. Modulation of NMDA receptor subunit mRNA in butorphanol-tolerant and -withdrawing rats. Neurochem Res. Dec 2000;25(12):1603–11.

17. Murray F, Harrison NJ, Grimwood S, Bristow LJ, Hutson PH. Nucleus accumbens NMDA receptor subunit expression and function is enhanced in morphine-dependent rats. Eur J Pharmacol. May 21 2007;562(3):191–7. doi:10.1016/j.ejphar.2007.01.027

18. Hopf FW. Do specific NMDA receptor subunits act as gateways for addictive behaviors? Genes Brain Behav. Jan 2017;16(1):118–138. doi:10.1111/gbb.12348

19. Koob GF, Volkow ND. Neurobiology of addiction: a neurocircuitry analysis. Lancet Psychiatry. Aug 2016;3(8):760–73. doi:10.1016/S2215-0366(16)00104-8

20. van Huijstee AN, Mansvelder HD. Glutamatergic synaptic plasticity in the mesocorticolimbic system in addiction. Front Cell Neurosci. 2014;8:466. doi:10.3389/fncel.2014.00466

21. Fluyau D, Revadigar N, Pierre CG. Clinical benefits and risks of N-Methyl-D-aspartate receptor antagonists to treat severe opioid use disorder: a systematic review. Drug and Alcohol Dependence. 2020/01/11/2020:107845. doi:https://doi.org/10.1016/j.drugalcdep.2020.107845

22. Xi ZX, Stein EA. Blockade of ionotropic glutamatergic transmission in the ventral tegmental area reduces heroin reinforcement in rat. Psychopharmacology (Berl). Nov 2002;164(2):144–50. doi:10.1007/s00213-002-1190-3

23. Trujillo KA, Akil H. Inhibition of morphine tolerance and dependence by the NMDA receptor antagonist MK-801. Science. Jan 4 1991;251(4989):85–7.

24. Bisaga A, Sullivan MA, Glass A, et al. A placebo-controlled trial of memantine as an adjunct to injectable extended-release naltrexone for opioid dependence. J Subst Abuse Treat. May-Jun 2014;46(5):546–52. doi:10.1016/j.jsat.2014.01.005

25. Liu Y, Lin D, Wu B, Zhou W. Ketamine abuse potential and use disorder. Brain Res Bull. Sep 2016;126(Pt 1):68-73. doi:10.1016/j.brainresbull.2016.05.016

26. Tomek SE, Lacrosse AL, Nemirovsky NE, Olive MF. NMDA Receptor Modulators in the Treatment of Drug Addiction. Pharmaceuticals (Basel). Feb 06 2013;6(2):251–68. doi:10.3390/ph6020251

27. Lu GY, Wu N, Zhang ZL, Ai J, Li J. Effects of D-cycloserine on extinction and reinstatement of morphine-induced conditioned place preference. Neurosci Lett. Oct 10 2011;503(3):196–9. doi:10.1016/j.neulet.2011.08.034

28. Myers KM, Carlezon WA, Jr. D-cycloserine facilitates extinction of naloxone-induced conditioned place aversion in morphine-dependent rats. Biol Psychiatry. Jan 1 2010;67(1):85–7. doi:10.1016/j.biopsych.2009.08.015

29. Myers KM, Carlezon WA, Jr. Extinction of drug- and withdrawal-paired cues in animal models: relevance to the treatment of addiction. Neurosci Biobehav Rev. Nov 2010;35(2):285–302. doi:10.1016/j.neubiorev.2010.01.011

30. Kosten TA, DeCaprio JL, Rosen MI. The severity of naloxone-precipitated opiate withdrawal is attenuated by felbamate, a possible glycine antagonist. Neuropsychopharmacology. Dec 1995;13(4):323–33. doi:10.1016/0893-133X(95)00065-L

31. Das RK, Kamboj SK. Maintaining clinical relevance: considerations for the future of research into D-cycloserine and cue exposure therapy for addiction. Biol Psychiatry. Dec 1 2012;72(11):e29-30; author reply e31-2. doi:10.1016/j.biopsych.2012.05.030

32. Hofmann SG. D-cycloserine for treating anxiety disorders: making good exposures better and bad exposures worse. Depress Anxiety. Mar 2014;31(3):175–7. doi:10.1002/da.22257

33. Haring R, Stanton PK, Scheideler MA, Moskal JR. Glycine-like modulation of N-methyl-D-aspartate receptors by a monoclonal antibody that enhances long-term potentiation. J Neurochem. Jul 1991;57(1):323–32.

34. Stanton PK, Potter PE, Aguilar J, Decandia M, Moskal JR. Neuroprotection by a novel NMDAR functional glycine site partial agonist, GLYX-13. Neuroreport. Aug 26 2009;20(13):1193–7. doi:10.1097/WNR.0b013e32832f5130

35. Moskal JR, Kuo AG, Weiss C, et al. GLYX-13: a monoclonal antibody-derived peptide that acts as an N-methyl-D-aspartate receptor modulator. Neuropharmacology. Dec 2005;49(7):1077–87. doi:10.1016/j.neuropharm.2005.06.006

36. Donello JE, Banerjee P, Li YX, et al. Positive N-Methyl-D-Aspartate Receptor Modulation by Rapastinel Promotes Rapid and Sustained Antidepressant-Like Effects. Int J Neuropsychopharmacol. Mar 1 2019;22(3):247–259. doi:10.1093/ijnp/pyy101

37. Moskal JR, Burgdorf JS, Stanton PK, et al. The Development of Rapastinel (Formerly GLYX-13); A Rapid Acting and Long Lasting Antidepressant. Curr Neuropharmacol. 2017;15(1):47–56.

38. Garay RP, Zarate CA, Jr., Charpeaud T, et al. Investigational drugs in recent clinical trials for treatment-resistant depression. Expert Rev Neurother. Jun 2017;17(6):593–609. doi:10.1080/14737175.2017.1283217

39. Newport DJ, Carpenter LL, McDonald WM, et al. Ketamine and Other NMDA Antagonists: Early Clinical Trials and Possible Mechanisms in Depression. Am J Psychiatry. Oct 2015;172(10):950–66. doi:10.1176/appi.ajp.2015.15040465

40. Zhang XL, Sullivan JA, Moskal JR, Stanton PK. A NMDA receptor glycine site partial agonist, GLYX-13, simultaneously enhances LTP and reduces LTD at Schaffer collateral-CA1 synapses in hippocampus. Neuropharmacology. Dec 2008;55(7):1238–50. doi:10.1016/j.neuropharm.2008.08.018

41. Burgdorf J, Zhang XL, Weiss C, et al. The long-lasting antidepressant effects of rapastinel (GLYX-13) are associated with a metaplasticity process in the medial prefrontal cortex and hippocampus. Neuroscience. Nov 12 2015;308:202-11. doi:10.1016/j.neuroscience.2015.09.004

42. Burgdorf J, Zhang XL, Weiss C, et al. The N-methyl-D-aspartate receptor modulator GLYX-13 enhances learning and memory, in young adult and learning impaired aging rats. Neurobiol Aging. Apr 2011;32(4):698–706. doi:10.1016/j.neurobiolaging.2009.04.012

43. Moskal JR, Burch R, Burgdorf JS, et al. GLYX-13, an NMDA receptor glycine site functional partial agonist enhances cognition and produces antidepressant effects without the psychotomimetic side effects of NMDA receptor antagonists. Expert Opin Investig Drugs. Feb 2014;23(2):243–54. doi:10.1517/13543784.2014.852536

44. Squeglia LM, Fadus MC, McClure EA, Tomko RL, Gray KM. Pharmacological Treatment of Youth Substance Use Disorders. J Child Adolesc Psychopharmacol. Aug 2019;29(7):559–572. doi:10.1089/cap.2019.0009

45. Villar VM, Bhargava HN. Pharmacodynamics and kinetics of loss of tolerance and physical dependence on morphine induced by pellet implantation in the rat. Pharmacology. 1992;45(6):319–28. doi:10.1159/000139017

46. Kosersky DS, Kowolenko MD, Howes JF. Physical dependence and tolerance development after chronic exposure to low levels of morphine in the rat. Pharmacol Biochem Behav. Apr 1980;12(4):625–8. doi:10.1016/0091-3057(80)90199-9

47. Yoburn BC, Chen J, Huang T, Inturrisi CE. Pharmacokinetics and pharmacodynamics of subcutaneous morphine pellets in the rat. J Pharmacol Exp Ther. Nov 1985;235(2):282–6.

48. Gellert VF, Holtzman SG. Development and maintenance of morphine tolerance and dependence in the rat by scheduled access to morphine drinking solutions. J Pharmacol Exp Ther. Jun 1978;205(3):536–46.

49. Burgdorf J, Zhang XL, Nicholson KL, et al. GLYX-13, a NMDA receptor glycine-site functional partial agonist, induces antidepressant-like effects without ketamine-like side effects. Neuropsychopharmacology. Apr 2013;38(5):729–42. doi:10.1038/npp.2012.246

50. Mueller D, Perdikaris D, Stewart J. Persistence and drug-induced reinstatement of a morphine-induced conditioned place preference. Behav Brain Res. Nov 15 2002;136(2):389–97.

51. Ghoreishi-Haack N, Priebe JM, Aguado JD, et al. NYX-2925 Is a Novel N-Methyl-d- Aspartate Receptor Modulator that Induces Rapid and Long-Lasting Analgesia in Rat Models of Neuropathic Pain. J Pharmacol Exp Ther. Sep 2018;366(3):485–497. doi:10.1124/jpet.118.249409

52. Khan MA, Houck DR, Gross AL, et al. NYX-2925 is a novel NMDA receptor-specific spirocyclic-beta-lactam that modulates synaptic plasticity processes associated with learning and memory. Int J Neuropsychopharmacol. Nov 1 2017;doi:10.1093/ijnp/pyx096

53. Kandasamy R, Lee AT, Morgan MM. Depression of home cage wheel running is an objective measure of spontaneous morphine withdrawal in rats with and without persistent pain. Pharmacol Biochem Behav. May 2017;156:10-15. doi:10.1016/j.pbb.2017.03.007

54. Cicero TJ, Nock B, Meyer ER. Gender-linked differences in the expression of physical dependence in the rat. Pharmacol Biochem Behav. Jun 2002;72(3):691–7. doi:10.1016/s0091-3057(02)00740-2

55. Kelsey JE, Aranow JS, Matthews RT. Context-specific morphine withdrawal in rats: duration and effects of clonidine. Behav Neurosci. Oct 1990;104(5):704–10. doi:10.1037//0735-7044.104.5.704

56. Craft RM, Stratmann JA, Bartok RE, Walpole TI, King SJ. Sex differences in development of morphine tolerance and dependence in the rat. Psychopharmacology (Berl). Mar 1999;143(1):1–7.

57. Towers EB, Tunstall BJ, McCracken ML, Vendruscolo LF, Koob GF. Male and female mice develop escalation of heroin intake and dependence following extended access. Neuropharmacology. Jun 2019;151:189-194. doi:10.1016/j.neuropharm.2019.03.019

58. McQueen K, Murphy-Oikonen J. Neonatal Abstinence Syndrome. N Engl J Med. Dec 22 2016;375(25):2468–2479. doi:10.1056/NEJMra1600879

59. Barr GA, Zmitrovich A, Hamowy AS, Liu PY, Wang S, Hutchings DE. Neonatal withdrawal following pre- and postnatal exposure to methadone in the rat. Pharmacol Biochem Behav. May 1998;60(1):97–104.

60. Ceger P, Kuhn CM. Opiate withdrawal in the neonatal rat: relationship to duration of treatment and naloxone dose. Psychopharmacology (Berl). Jun 2000;150(3):253–9.

61. Barr GA, McPhie-Lalmansingh A, Perez J, Riley M. Changing mechanisms of opiate tolerance and withdrawal during early development: animal models of the human experience. ILAR J. 2011;52(3):329–41. doi:10.1093/ilar.52.3.329

62. Windh RT, Little PJ, Kuhn CM. The ontogeny of mu opiate tolerance and dependence in the rat: antinociceptive and biochemical studies. J Pharmacol Exp Ther. Jun 1995;273(3):1361–74.

63. Jimenez-Romero F, Bis-Humbert C, Garcia-Fuster MJ. Adolescent morphine induces emotional signs of withdrawal paired with neurotoxicity selectively in male rats: Female resilience. Neurosci Lett. Jan 10 2020;715:134625. doi:10.1016/j.neulet.2019.134625

64. Doherty JM, Frantz KJ. Attenuated effects of experimenter-administered heroin in adolescent vs. adult male rats: physical withdrawal and locomotor sensitization. Psychopharmacology (Berl). Feb 2013;225(3):595–604. doi:10.1007/s00213-012-2847-1

65. Karami M, Zarrindast MR. Morphine sex-dependently induced place conditioning in adult Wistar rats. Eur J Pharmacol. Mar 17 2008;582(1-3):78-87. doi:10.1016/j.ejphar.2007.12.010

66. Cicero TJ, Ennis T, Ogden J, Meyer ER. Gender differences in the reinforcing properties of morphine. Pharmacol Biochem Behav. Jan 1 2000;65(1):91–6. doi:10.1016/s0091-3057(99)00174-4

67. Timar J, Sobor M, Kiraly KP, et al. Peri, pre and postnatal morphine exposure: exposure-induced effects and sex differences in the behavioural consequences in rat offspring. Behav Pharmacol. Feb 2010;21(1):58–68. doi:10.1097/FBP.0b013e3283359f39

68. Lynch WJ, Carroll ME. Sex differences in the acquisition of intravenously self-administered cocaine and heroin in rats. Psychopharmacology (Berl). May 1999;144(1):77–82.

69. Cicero TJ, Aylward SC, Meyer ER. Gender differences in the intravenous self-administration of mu opiate agonists. Pharmacol Biochem Behav. Feb 2003;74(3):541–9.

70. Hempel BJ, Crissman ME, Imanalieva A, et al. Cross-Generational THC Exposure Alters Heroin Reinforcement in Adult Male Offspring. Drug Alcohol Depend. Jul 1 2020;212:107985. doi:10.1016/j.drugalcdep.2020.107985

71. Venniro M, Zhang M, Shaham Y, Caprioli D. Incubation of Methamphetamine but not Heroin Craving After Voluntary Abstinence in Male and Female Rats. Neuropsychopharmacology. Apr 2017;42(5):1126–1135. doi:10.1038/npp.2016.287

72. Caille S, Espejo EF, Reneric JP, Cador M, Koob GF, Stinus L. Total neurochemical lesion of noradrenergic neurons of the locus ceruleus does not alter either naloxone-precipitated or spontaneous opiate withdrawal nor does it influence ability of clonidine to reverse opiate withdrawal. J Pharmacol Exp Ther. Aug 1999;290(2):881–92.

73. Frenois F, Cador M, Caille S, Stinus L, Le Moine C. Neural correlates of the motivational and somatic components of naloxone-precipitated morphine withdrawal. Eur J Neurosci. Oct 2002;16(7):1377–89.

74. Nestler EJ. Reflections on: “A general role for adaptations in G-Proteins and the cyclic AMP system in mediating the chronic actions of morphine and cocaine on neuronal function”. Brain Res. Aug 15 2016;1645:71-4. doi:10.1016/j.brainres.2015.12.039

75. Rasmussen K, Beitner-Johnson DB, Krystal JH, Aghajanian GK, Nestler EJ. Opiate withdrawal and the rat locus coeruleus: behavioral, electrophysiological, and biochemical correlates. J Neurosci. Jul 1990;10(7):2308–17.

76. Rasmussen K. The role of the locus coeruleus and N-methyl-D-aspartic acid (NMDA) and AMPA receptors in opiate withdrawal. Neuropsychopharmacology. Dec 1995;13(4):295–300. doi:10.1016/0893-133X(95)00082-O

77. Rasmussen K, Brodsky M, Inturrisi CE. NMDA antagonists and clonidine block c-fos expression during morphine withdrawal. Synapse. May 1995;20(1):68–74. doi:10.1002/syn.890200110

78. Rasmussen K, Fuller RW, Stockton ME, Perry KW, Swinford RM, Ornstein PL. NMDA receptor antagonists suppress behaviors but not norepinephrine turnover or locus coeruleus unit activity induced by opiate withdrawal. Eur J Pharmacol. May 2 1991;197(1):9–16. doi:10.1016/0014-2999(91)90358-w

79. Kato T, Duman RS. Rapastinel, a novel glutamatergic agent with ketamine-like antidepressant actions: Convergent mechanisms. Pharmacol Biochem Behav. Jan 2020;188:172827. doi:10.1016/j.pbb.2019.172827

80. Aguilar MA, Rodriguez-Arias M, Minarro J. Neurobiological mechanisms of the reinstatement of drug-conditioned place preference. Brain Res Rev. Mar 2009;59(2):253–77. doi:10.1016/j.brainresrev.2008.08.002

81. Popik P, Wrobel M, Bisaga A. Reinstatement of morphine-conditioned reward is blocked by memantine. Neuropsychopharmacology. Jan 2006;31(1):160–70. doi:10.1038/sj.npp.1300760

82. Ribeiro Do Couto B, Aguilar MA, Manzanedo C, Rodriguez-Arias M, Minarro J. NMDA glutamate but not dopamine antagonists blocks drug-induced reinstatement of morphine place preference. Brain Res Bull. Jan 30 2005;64(6):493–503. doi:10.1016/j.brainresbull.2004.10.005

83. Girardi BA, Fabbrin S, Wendel AL, Mello CF, Rubin MA. Spermidine, a positive modulator of the NMDA receptor, facilitates extinction and prevents the reinstatement of morphine-induced conditioned place preference in mice. Psychopharmacology (Berl). Mar 2020;237(3):681–693. doi:10.1007/s00213-019-05403-z

84. Fasipe OJ. The emergence of new antidepressants for clinical use: Agomelatine paradox versus other novel agents. IBRO Rep. Jun 2019;6:95-110. doi:10.1016/j.ibror.2019.01.001

85. Wilkinson ST, Sanacora G. A new generation of antidepressants: an update on the pharmaceutical pipeline for novel and rapid-acting therapeutics in mood disorders based on glutamate/GABA neurotransmitter systems. Drug Discov Today. Feb 2019;24(2):606–615. doi:10.1016/j.drudis.2018.11.007

86. Koob GF. Neurobiology of Opioid Addiction: Opponent Process, Hyperkatifeia, and Negative Reinforcement. Biol Psychiatry. Jan 1 2020;87(1):44–53. doi:10.1016/j.biopsych.2019.05.023

87. Kenny PJ, Hoyer D, Koob GF. Animal Models of Addiction and Neuropsychiatric Disorders and Their Role in Drug Discovery: Honoring the Legacy of Athina Markou. Biol Psychiatry. Jun 1 2018;83(11):940–946. doi:10.1016/j.biopsych.2018.02.009

